# Governance scale and network structure shape pollinator recovery under pesticide reduction

**DOI:** 10.64898/2026.05.26.728062

**Authors:** Adrija Datta, Arnob Ray, Udit Bhatia

## Abstract

Reducing pesticide risks while maintaining food production remains a central challenge for sustainable agriculture. Although pesticide reduction is pursued through centralized regulation and farm-level Integrated Pest Management, how these governance pathways translate into pollinator recovery in agroecological systems remains poorly understood. Existing ecological network models often treat pesticide pressure asm external forcing and management actions as fixed parameters, limiting their ability to capture feedbacks among governance decisions, network structure, and population dynamics. Here, we develop a dynamical framework that embeds pesticide management within tripartite pollinator-plant-pest networks using a policy variable and a farm-level adoption variable. Across empirical and synthetic networks, we show that recovery is not determined by pesticide reduction alone, but by how management acts through ecological interaction structure. More modular networks require stronger intervention, and pollinators with similar degrees show different recovery outcomes, indicating that degree alone does not determine recovery potential. Further, increasing policy strength generally expands the persistence domain more than increasing farmer adoption alone. These results show that pesticide reduction does not automatically yield ecological recovery, and effective strategies must match governance scale to ecological condition and network structure.

## 1 Introduction

Biodiversity governance is increasingly focused on sustaining ecosystem functions and contributions of nature to people within human dominated landscapes, particularly agriculture. A prominent example is the Kunming Montreal Global Biodiversity Framework adopted under the Convention on Biological Diversity, which calls for reducing the overall risk of pesticides and highly hazardous chemicals by at least 50% by 2030 [1]. This emphasis reflects the growing recognition that agricultural intensification and land use change, including the conversion of forests and semi natural habitats to croplands, simplify landscapes and erode insect diversity and abundance [2–4]. At the same time, intensive reliance on synthetic pesticides and herbicides can disrupt ecological interactions and generate environmental and human health risks across large spatial scales [5–12]. Chemical inputs may stabilize yields and farm income in the short term, but can also harm non target beneficial insects, including pollinators and natural enemies of pests [13], thus weakening the ecological processes that support resilient food production.

Agricultural ecosystems are structured by interacting mutualistic and antagonistic processes that jointly determine productivity and ecological stability. Mutualistic interactions, particularly pollination, enhance fruit and seed production and can stimulate plant population growth, whereas antagonistic interactions such as herbivory by insect pests suppress plant performance and may destabilize agroecosystems during outbreaks. Managing the balance between these opposing forces is therefore central to reconciling biodiversity conservation with food production. Two broad governance pathways are commonly proposed to reduce the ecological risks associated with pesticide use. The first is centralized policy regulation that constrains chemical inputs through bans, restrictions, or risk based standards [14]. The second is decentralized Integrated Pest Management, abbreviated as IPM [15, 16], which combines biological, cultural, and selective chemical controls to suppress pests while minimizing impacts on beneficial insects. Despite their prominence in both policy and agricultural practice, the ecological consequences of these governance pathways remain difficult to evaluate at the system level because pesticide impacts propagate through networks of interacting species rather than acting on isolated populations.

Network approaches provide a natural framework for understanding how ecological responses to disturbance propagate through interacting species. Early empirical studies of agroecosystems showed that species are embedded in networks spanning multiple interaction types, including pollination, herbivory, and parasitism, and that these subnetworks can differ in their robustness to species loss [17]. Subsequent work formalized this perspective within multilayer and multiplex ecological network theory, demonstrating that coupling among interaction layers can generate stability and collapse dynamics that differ qualitatively from those predicted by single layer networks [18]. Despite these advances, much of the dynamical network literature still treats mutualistic and antagonistic interactions separately or within simplified bipartite representations [19–24]. Empirical and theoretical studies increasingly show that ecosystem stability depends on the balance among interaction types rather than on any single layer alone [25]. Dynamical pollinator plant herbivore models indicate that pollination can promote coexistence but may also destabilize dynamics, whereas herbivory suppresses growth while sometimes contributing to stability, implying that persistence emerges from the interaction of these opposing forces [26]. Studies explicitly combining mutualistic and antagonistic interactions further suggest that coupled layers can generate abrupt transitions and multiple critical points as interaction composition changes [27–31].

A parallel line of research has begun to incorporate pesticides as an explicit stressor in ecological network dynamics. These studies show that contamination can induce bistability and abrupt collapse in multispecies plant pollinator communities and that targeted ecological interventions may delay tipping points [32]. However, in most existing models management actions themselves, including pesticide regulation and IPM adoption, are treated as externally imposed parameters rather than as endogenous controls coupled to network dynamics. Anthropogenic stressors are typically represented as fixed forcing terms and evaluated without feedbacks among management decisions, ecological network structure, and population dynamics [32–36]. Consequently, it remains unclear how alternative governance pathways reshape the recoverability of ecological networks once systems have been pushed toward collapse and how that recoverability depends on interaction topology.

Against this background, we address two questions. First, how can pesticide management, capturing both policy regulation and farm-level adoption of IPM, be incorporated into dynamical models of pollinator persistence and recovery in coupled pollinator-plant-pest systems? Second, how do these management interventions interact with ecological network topology to shape recovery thresholds in agroecosystems? To answer these questions, we develop a dynamical framework that embeds pesticide pressure directly in tripartite ecological dynamics and represents governance through centralized regulation and decentralized adoption of reduced-pesticide practices. We then analyze recovery across empirical and synthetic networks to examine how initial ecological conditions, interaction structure, and governance scale jointly shape recoverability. In this way, the framework links pesticide risk reduction to ecological recovery in a systems-level setting.

## 2 Results

To investigate the impact of pesticide management on pollinator persistence in agroecosystems, we study a dynamical model (Eq. (1) in Sec. 4.2) that captures coupled pollinator–plant–pest interactions, along with the recovery of pollinator populations from extinction. In this framework, pesticide application suppresses pest populations while simultaneously increasing pollinator mortality, thereby weakening pollination benefits to plants. The tripartite interaction structure, ecological feedback pathways, and coupled governance controls are illustrated schematically in Fig. 1a–d. Governance interventions enter the model through two control parameters which are the adoption of integrated pest management by farmers (*x*_*u*_) and the strength of the centralized policy limiting pesticide use (*f*). Together, these parameters determine the effective pesticide pressure experienced by ecological communities.

**Figure 1:**
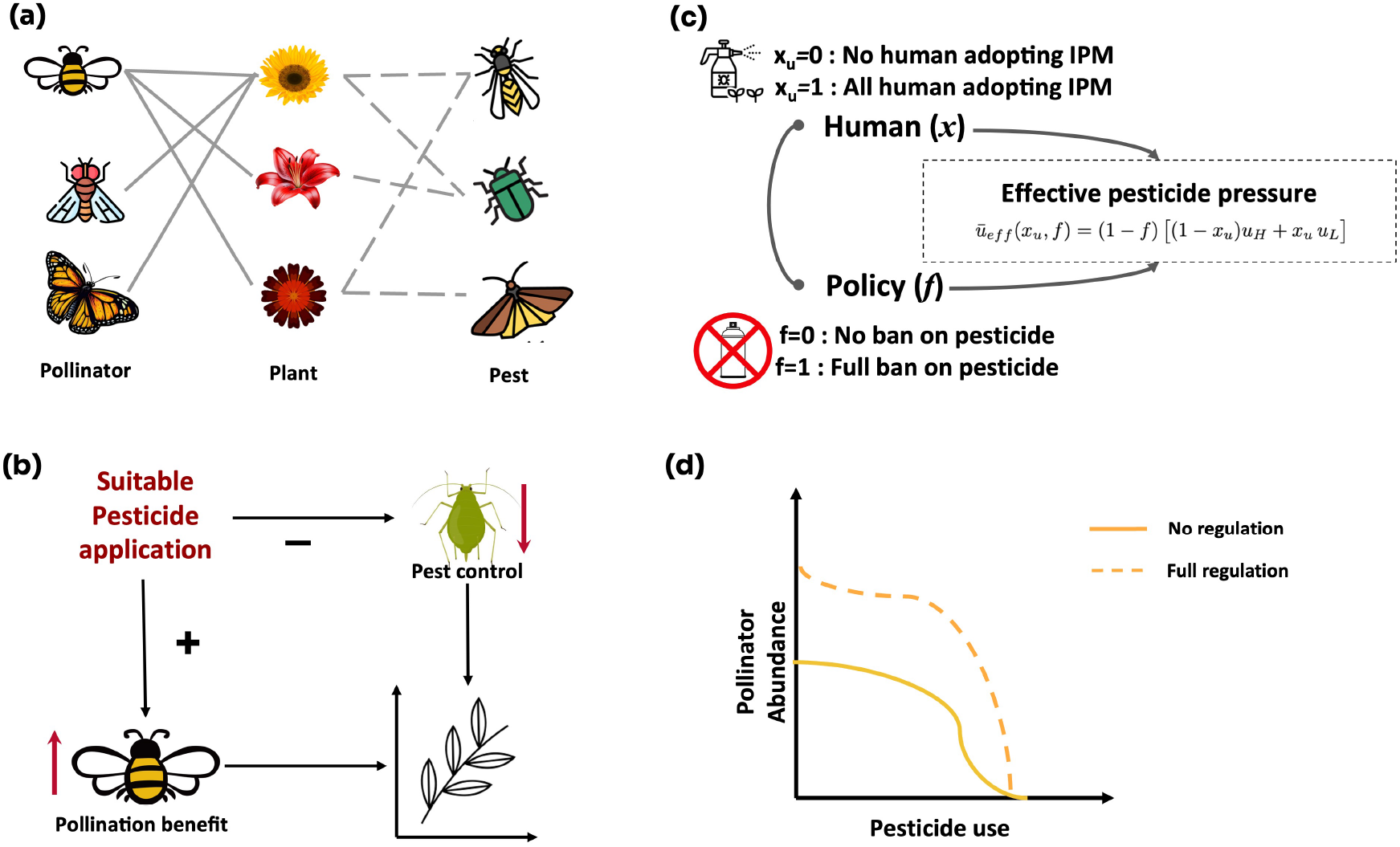
Conceptual framework linking pollination, pest control, and pesticide regulation. (a) Schematic representation of the tripartite pollinator–plant–pest network, illustrating mutualistic (solid lines) and antagonistic (dashed lines) interactions. (b) Conceptual feedback pathways showing how management of pesticide application simultaneously suppresses pests and increases pollination services. (c) Coupled human-policy system defining effective pesticide pressure as a function of IPM adoption (*x*_*u*_) and regulatory policy strength (*f*). (d) Conceptual response of pollinator abundance to increasing pesticide use under unregulated (solid line) and fully regulated (dashed line) scenarios.

### 2.1 Topological drivers of pollinator recovery in pollinator-plant-pest networks

We implement the model (Eq. 1) in an empirical tripartite ecological network (Sec. 4.1) compiled from Shinohara et al. [37], which simultaneously reports mutualistic pollinator-plant interactions and antagonistic plant-pest interactions. The network contains 23 pollinator species, 36 plant species, and 51 pest species (Fig. 2a). To examine how pesticide management affects ecological recovery, we vary the aforementioned control parameters (*x*_*u*_ and *f*) between 0 and 1 as they govern the pressure of pesticides. Increasing the intensity of management parameters induces recovery transitions for subsets of pollinator species, shifting the system from an extinction regime to a persistence regime (Fig. 2b). These transitions may occur gradually or abruptly, reflecting nonlinear feedbacks between ecological interactions and pesticide mortality.

**Figure 2:**
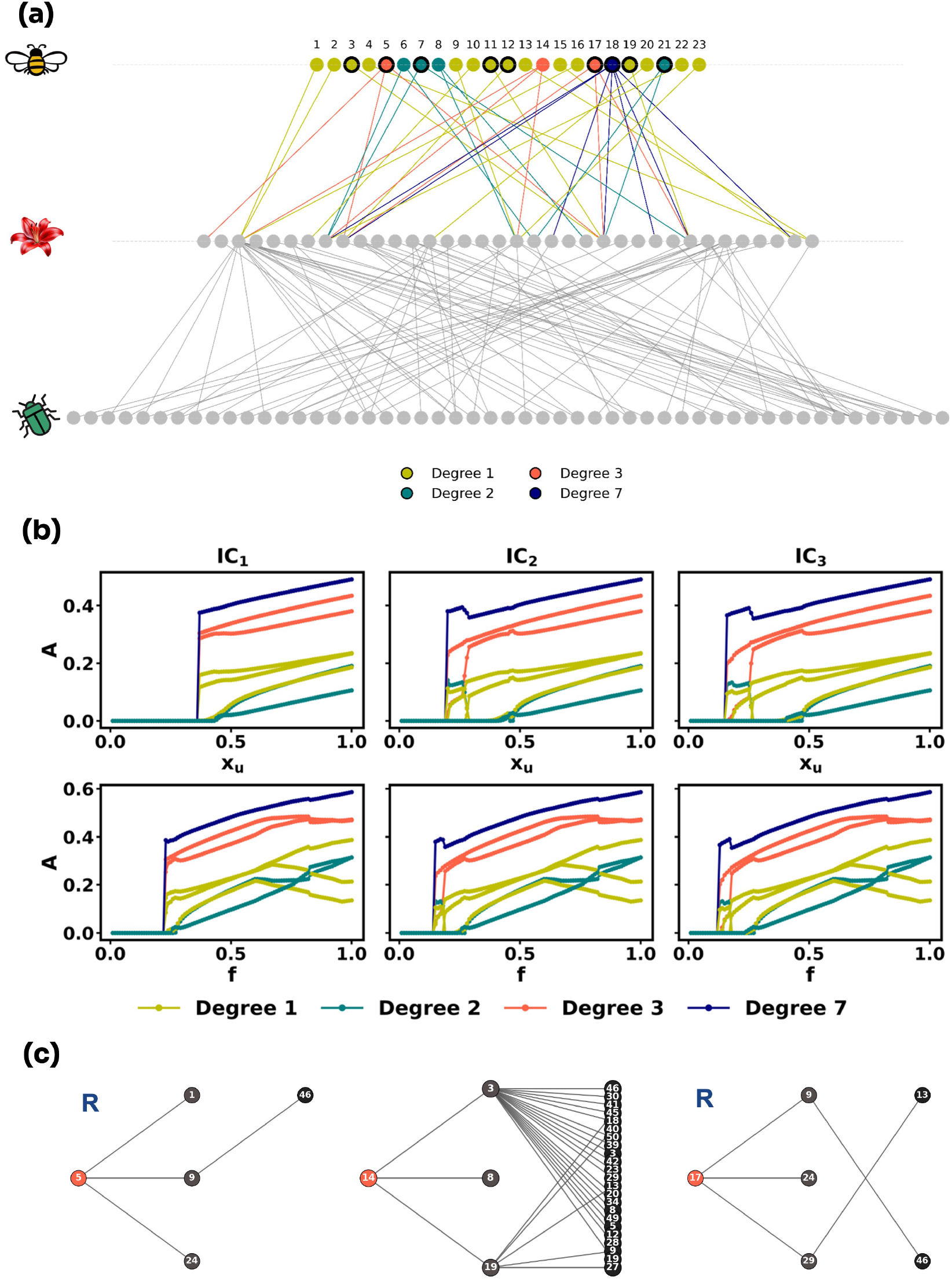
Dynamics of pollinator–plant–pest interactions and recovery. (a) Tripartite interaction network showing pollinators (top panel), plants (middle panel), and pests (bottom). Pollinator nodes are colored by their degree, while those outlined in black highlight the subset of species that undergo a transition from an extinct state to recovery, as shown in panel (b). (b) Transitions of pollinators from extinction to persistence under varying pesticide-management parameters across three initial conditions (IC_1_, IC_2_, and IC_3_). The upper panel fixes policy strength at *f* = 0.1, while the lower panel fixes IPM adoption at *x*_*u*_ = 0.1. (c) Representative degree-3 pollinator motifs illustrating contrasting levels of indirect pest exposure and corresponding recovery outcomes. Annotations indicate recovery status (R), with connections to pests occurring via shared plant partners, as visible in the figure.

Recovery patterns are strongly dependent on the structure of the network. Although some pollinator species recover as the intensity of management increases, others remain in extinct state even when they have identical degrees (Fig. 2c and Sec. B in Supporting Information). This indicates that degree alone does not determine recovery potential. Instead, tripartite network structure plays an important role (Sec. B.1 in Supporting iInformation). Examination of local interaction motifs reveals that pollinators connected to plants strongly associated with pest species remain vulnerable to collapse. In such motifs, pesticide-driven changes in pest abundance alter plant dynamics and cascade back to pollinators, preventing recovery even under improved management conditions (Fig. 2c).

Recovery dynamics also depend on the initial ecological state. To illustrate this effect, we assign identical initial abundances to all species within each guild. For all simulations, species are initialized uniformly, so that 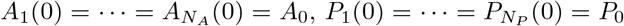, and 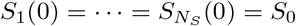 The three initial conditions used in Fig. 2 are

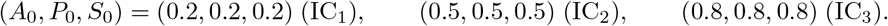

For low initial abundances (IC_1_), pollinator recovery occurs only beyond a relatively high IPM threshold 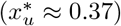 when the policy factor is fixed at *f* = 0.1. In contrast, for IC_2_ and IC_3_, recovery occurs at lower IPM values 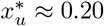 and 0.17, respectively). Similarly, when IPM adoption is fixed at *x*_*u*_ = 0.1, the policy threshold (*f*^*^) required for recovery decreases as initial pollinator abundance increases (Fig. 2b). These results indicate that when ecological populations begin at low abundance, stronger management interventions are required to restore pollinator persistence.

To generalize the results across different tripartite network properties, we tested our hypothesis across a set of 30 synthetic pollinator–plant–pest networks with identical species richness but varying modularity (Sec. 4.4), recovery dynamics also show strong dependence on network architecture. Increasing modularity systematically raises the critical management thresholds required for recovery and reduces the fraction of pollinator species that successfully transition to persistence (Sec. B.2 in Supporting Information). These results indicate that more compartmentalized ecological networks require stronger interventions to achieve system-wide recovery.

### 2.2 Quantifying resilience of pollinators under initial-condition sensitivity

Figure 3a shows the basin of attraction of the pollinator of the highest-degree (*A*_18_) for persistent and non-persistent states across the (*x*_*u*_, *f*) control plane and reveals a strong sensitivity of recovery outcomes to management intensity. We quantify these dynamics using basin stability (BS), defined as the fraction of initial states that converge to the attractor (Sec. 4.5). At low levels of both IPM adoption and policy intervention (e.g., *x*_*u*_ = 0.1, *f* = 0.1), the persistence basin is nearly absent, with basin stability values approaching zero, indicating that almost all initial ecological states collapse to extinction. As management intensity increases, even modestly, the persistence basin expands rapidly. For example, at *f* = 0.1, basin stability increases from approximately 0.0 at *x*_*u*_ = 0.1 to about 0.395 at *x*_*u*_ = 0.2, and further to 0.698 at *x*_*u*_ = 0.3, showing a strong nonlinear response to changes in farmer behavior.

**Figure 3:**
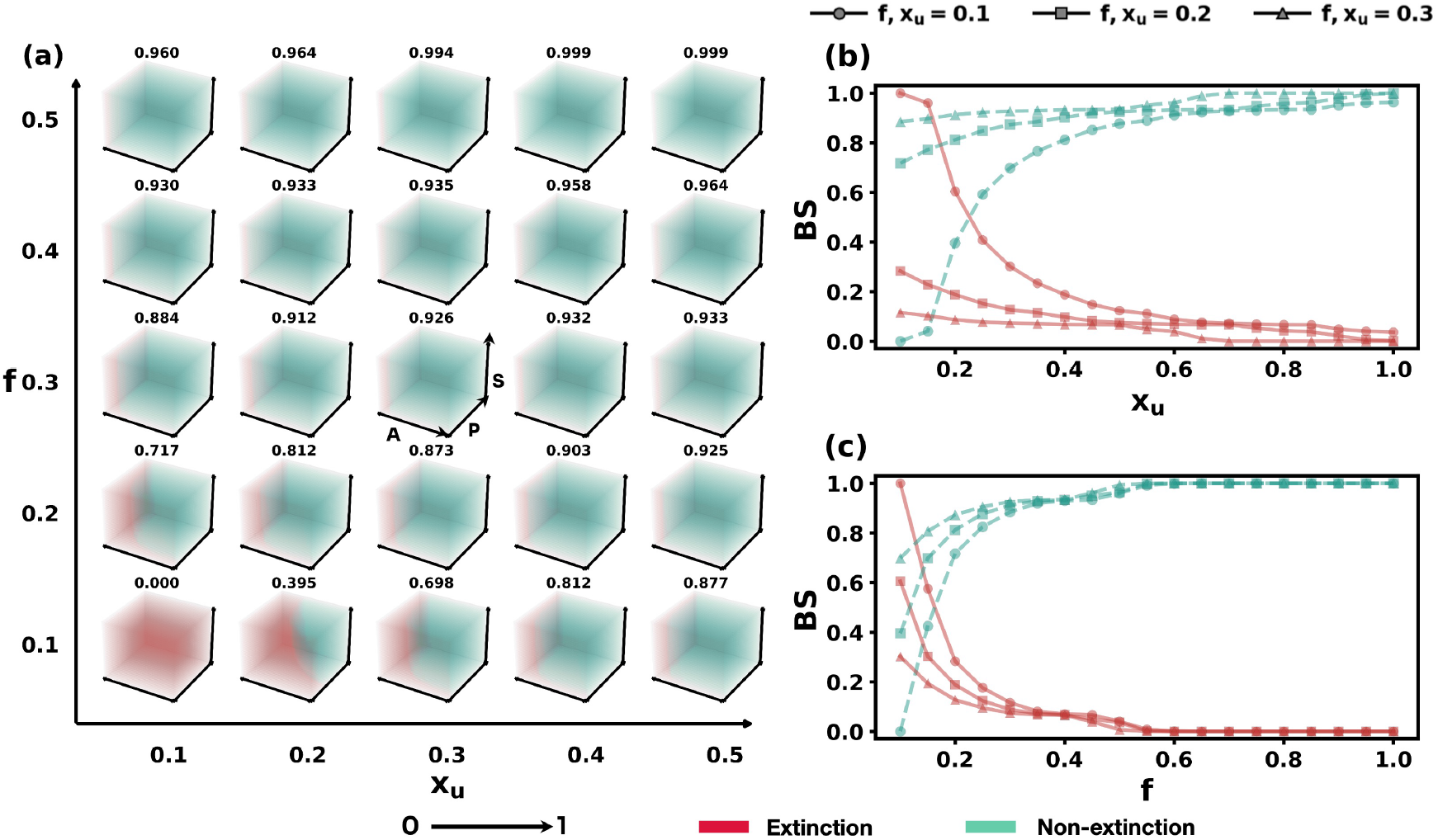
Initial-condition dependence of recovery outcomes for the highest-degree pollinator. (a) Basin of attraction in the (*x*_*u*_, *f*) parameter space, showing the persistence probability (basin stability of non-extinction state) of *A*_18_ across management combinations. (b) Basin stability as a function of *x*_*u*_ for fixed *f*. (c) Basin stability as a function of *f* for fixed *x*_*u*_. Basin of attraction and basin stability of the extinct state are shown in red, while those of the non-extinct state are shown in green.

This expansion becomes more pronounced under stronger policy regimes. At *f* = 0.3, basin stability values already exceed 0.88 at *x*_*u*_ = 0.1, reaching 0.926 at *x*_*u*_ = 0.3 and exceeding 0.93 beyond *x*_*u*_ = 0.4. Under even higher policy strength (*f* ≥ 0.4), the persistence basin dominates the phase space, with basin stability consistently above 0.93 and approaching 0.99 for larger *x*_*u*_, indicating near-certain recovery regardless of initial ecological conditions. These patterns demonstrate that increasing either IPM adoption or policy strength shifts the system away from extinction-dominated dynamics toward persistence-dominated regimes.

Consistent with this geometry of basin of attraction, varying initial ecological conditions leads to shifts in the bifurcation points associated with pollinator recovery in Fig. 2b. These shifts arise because the system exhibits bistability, with coexisting attractors corresponding to extinction (near-zero pollinator abundance) and persistence (positive pollinator populations). When initial conditions are within the extinction basin, recovery requires crossing a critical threshold in either *x*_*u*_ or *f*, whereas initial states within the persistence basin recover without additional intervention.

BS increases monotonically with both management parameters (Fig. 3b,c), but the rate of increase differs substantially between them. For instance, in Fig. 3b, increasing *x*_*u*_ from 0.1 to 0.4 at low policy levels results in only gradual increases in BS, whereas at higher *f*, the same increase in *x*_*u*_ rapidly pushes BS toward unity. In contrast, Fig. 3c shows that increasing *f* leads to a much sharper transition. BS rises from near zero to above 0.9 within a relatively narrow range of *f*, even for low *x*_*u*_.

This asymmetry indicates that policy-driven reductions in pesticide pressure more effectively reshape the global stability landscape of the system than decentralized IPM adoption alone. While increasing *x*_*u*_ incrementally enlarges the persistence basin, increasing *f* produces a more rapid and system-wide expansion, effectively eliminating the extinction basin over a broad region of parameter space. Consequently, the probability that ecological communities recover from low-abundance states increases much more efficiently under stronger policy intervention than under comparable increases in farm-level adoption.

### 2.3 Interplay of policy, farmer behaviour, and species interaction strength

To assess how mutualistic (*γ*_0_) and antagonistic (*ξ*_0_) coupling strengths shape pollinator recovery, we analyze the transition boundary separating extinction from persistence for the pollinator of the highest-degree (*A*_18_) for a fixed set of initial conditions ((*A*_0_, *P*_0_, *S*_0_) = (0.2, 0.2, 0.2) (IC_1_)). We compute the critical values of *x*_*u*_ 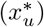 at fixed *f* = 0.1, 0.2, 0.3 (Fig. 4a–c) and of *f* (*f*^*^) at fixed *x*_*u*_ = 0.1, 0.2, 0.3 (Fig. 4d–f) across the *γ*_0_–*ξ*_0_ parameter space.

**Figure 4:**
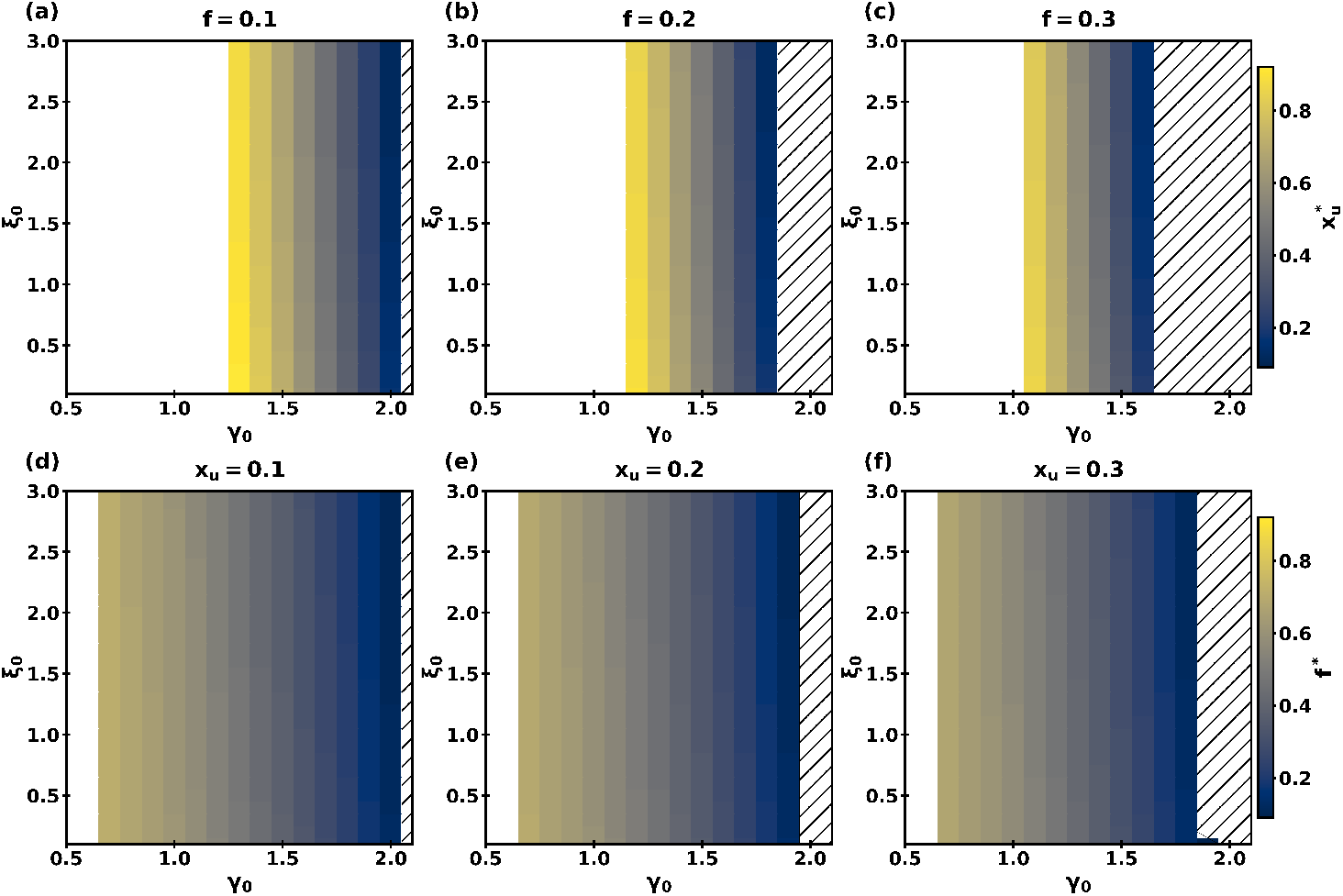
Influence of mutualistic and antagonistic interaction strengths on the transition point of management parameters governing the recovery of the highest degree pollinator (*A*_18_). Top panel shows critical adoption thresholds 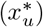 at fixed policy levels, and bottom panel shows critical policy thresholds (*f*^*^) at fixed adoption levels. Masked regions indicate the absence of a transition point within the scanned parameter range. White regions denote no recovery transition within the scanned range, whereas hatched regions denote parameter combinations for which persistence is maintained across the scanned range and no threshold is required. This is shown for IC_1_.

Recovery transitions are primarily governed by mutualistic coupling strength *γ*_0_. Across all panels, the transition boundary is nearly vertical, showing that recovery depends strongly on *γ*_0_ and only weakly on *ξ*_0_. As *γ*_0_ increases, the threshold of management required for recovery declines markedly. For example, for fixed *f* (Fig. 4a–c), at low *γ*_0_ (around 1.1–1.3), recovery requires high intervention (*x*_*u*_ ≳ 0.8), whereas at moderate *γ*_0_ (around 1.4–1.6), much lower levels (~ 0.3) are sufficient. In contrast, changes in antagonistic strength *ξ*_0_ shift the boundary only slightly, indicating that antagonistic interactions influence but do not control recovery outcomes. At higher mutualistic strengths (*γ*_0_ ≳ 1.6–1.7), the system enters a regime of permanent persistence when *f* is kept at higher value greater than 0.3, where pollinator populations remain stable across the full range of management conditions (*x*_*u*_ ∈ [0, 1]). This reflects a self-sustaining tripartite network in which strong plant–pollinator feedbacks buffer the system against external stress.

Comparing management pathways reveals consistent differences between IPM adoption (*x*_*u*_) and policy regulation (*f*). For similar ecological conditions, recovery typically requires higher levels of adoption than policy. For instance, at intermediate *γ*_0_ (around 1.3–1.5), recovery can occur at fixed policy levels (*f*^*^) of approximately 0.3–0.4, while adoption 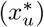 may need to reach 0.5–0.7. This suggests that policy reduces pesticide pressure more effectively at the system level, whereas adoption acts more gradually and depends more on ecological support. As policy strength increases (from *f* = 0.1 to *f* = 0.3), the bistable region becomes narrower, indicating that stronger regulation accelerates recovery and reduces sensitivity to pesticide management practices.

These results show that recovery in pollinator networks is primarily controlled by the strength of mutualistic interactions, with management acting to shift the system across this boundary. Strong ecological feedbacks lower the need for intervention, while policy-driven reductions in pesticide pressure more effectively stabilize recovery dynamics and expand the conditions under which persistence can be achieved.

## 3 Discussion

Pollinator recovery in agricultural ecosystems depends on how pesticide reduction propagates through coupled ecological and governance feedbacks. By embedding pesticide use, farmer behavior, and regulatory control in a tripartite pollinator-plant-pest model, we show that recoverability is shaped not only by the magnitude of intervention but also by the initial ecological state and the structure of species interactions. Communities beginning from depleted abundances require stronger intervention to return to persistence, and more compartmentalized networks are harder to restore.

In our parameterization, increasing policy strength typically expands the basin of persistence more than increasing IPM adoption alone. The results highlight that broad reductions in exposure can shift recovery conditions more rapidly than partial local uptake when systems are already close to collapse although the two controls are not structurally symmetric in the model, because policy rescales total pesticide use whereas adoption redistributes effort between conventional and low-pesticide management. Part of the stronger policy response therefore reflects how governance enters the effective pesticide-pressure term, not only ecological dynamics. Regulatory measures such as bans on highly toxic insecticides, restrictions on application during flowering, and landscape-scale buffer zones can reduce exposure across fields and habitats [38, 39], whereas IPM can reduce inputs while depending more strongly on uptake, knowledge, and local resource conditions [40, 41].

These findings argue against evaluating pesticide governance through isolated species responses alone. Recovery depends on when intervention occurs, how broadly it is applied, and how ecological interactions transmit its effects through the network. Local motifs and modular organization can trap recovery within parts of the community, while stronger mutualistic feedbacks lower the intervention needed for persistence. In that sense, governance scale and ecological structure jointly determine whether pesticide reduction translates into recovery.

Our work has certain limitations. Pesticide effects are represented through aggregate mortality terms rather than species-specific toxicological response functions, and policy regulation and IPM adoption are introduced as control parameters rather than empirical decision processes. The diversity of IPM practices, including biological control, habitat management, and cultural methods such as mulching, is collapsed into a single management dimension [42, 43]. The framework also omits implementation delays, spatial heterogeneity in exposure, explicit economic costs, and adaptive farmer behavior. Future work can relax these assumptions by coupling governance more realistically, incorporating spatial tripartite networks, and evaluating optimized management portfolios [44, 45].

Despite these limitations, the framework provides a systems-level way to connect pesticide reduction to ecological recovery in interacting communities. By linking governance decisions to tripartite ecological dynamics, it shows that recovery depends not only on how much pesticide pressure is reduced, but also on initial ecological condition, network structure, and the scale at which intervention acts. In that sense, the study offers a basis for designing pesticide reduction strategies that are better matched to ecological context rather than applied uniformly.

## 4 Materials and Methods

### 4.1 Empirical tripartite network

We use the empirical tripartite network reported by Shinohara et al. [37], which contains both pollinator-plant mutualistic interactions and plant-pest antagonistic interactions in the same agroecosystem setting. To maintain relevance to pesticide-induced changes in pollinator abundance, we have further restricted our selection to networks originating from agricultural or agro-ecosystem contexts. In particular, we have focused on systems situated in landscapes containing seminatural grasslands adjacent to croplands. These seminatural habitats are former rice fields where cultivation had been abandoned for at least ten years, and both the fields and their edges have remained unmowed. Such settings provide ecologically realistic conditions where pollinators, plants, and pests cooccur and interact under management regimes relevant to pesticide exposure. The network contains *N*_*A*_ = 23 pollinator species, *N*_*P*_ = 36 plant species, and *N*_*S*_ = 51 pest species (Fig. 2a). The pollinator–plant and plant–pest interactions are encoded as binary adjacency matrices, with the dynamical model assigning the corresponding interaction strengths.

### 4.2 Pollinator-plant-pest-pesticide (P4) model

We model the abundances of pollinators (*A*), plants (*P*), and pests (*S*) with a coupled nonlinear system that combines density regulation, mutualistic gains, antagonistic losses, and pesticide-induced mortality. Let *A*_*i*_(*t*), *P*_*i*_(*t*), and *S*_*i*_(*t*) denote the abundances of pollinator, plant, and pest species, respectively. The dynamics evolve in an (*N*_*A*_ + *N*_*P*_ + *N*_*S*_)-dimensional state space according to

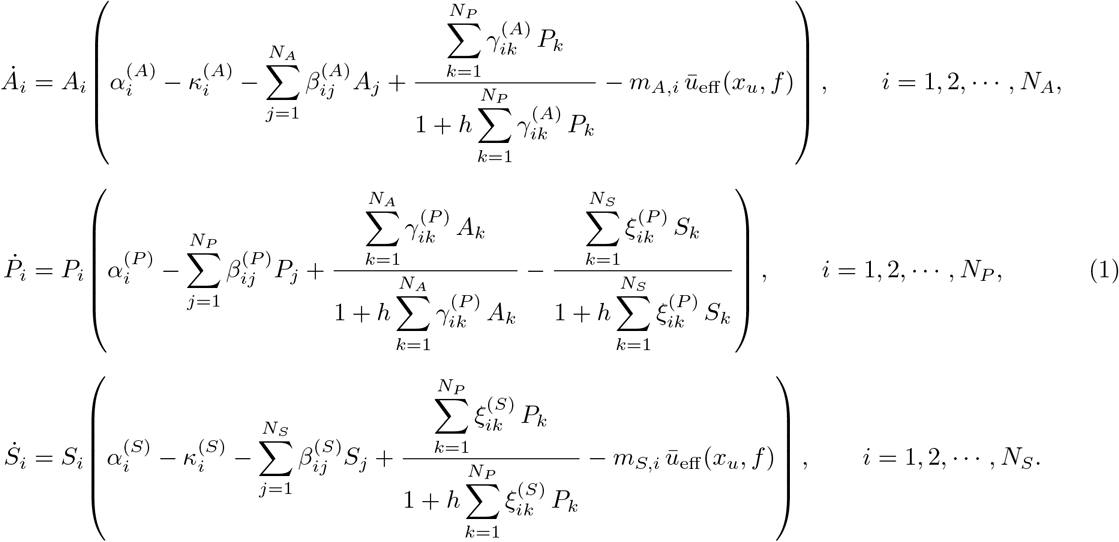

The pollinator and pest equations contain intrinsic growth, background mortality, intra- and interspecific competition, saturating gains from plant partners, and direct pesticide mortality. The plant equation contains intrinsic growth, competition, saturating mutualistic benefits from pollinators, and saturating losses to pests. Direct pesticide mortality acts only on pollinators and pests, reflecting non-target and target pesticide effects, respectively. Saturating interaction terms follow Holling type II responses [33, 46, 47].

### 4.3 Management controls and interaction parameterization

Governance enters the model through two control parameters. The policy factor *f* ∈ [0, 1] represents a reduction in pesticide use by implementing centralized restrictions on pesticide use, with *f* = 0 corresponding to the status quo and *f* = 1 to complete elimination of pesticide application. The adoption factor *x*_*u*_ ∈ [0, 1] denotes the fraction of farmers using low-pesticide or integrated pest management (IPM) practices. The mean pesticide intensity associated with farmer behaviour is

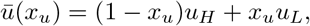

where *u*_*H*_ and *u*_*L*_ are the pesticide intensities associated with conventional and IPM management, respectively. Policy and adoption combine multiplicatively to define the effective pesticide pressure

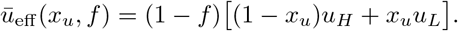

This effective pressure enters Eq. (1) as the pesticide-mortality term for pollinators and pests.

However, the comparison between policy and adoption must be interpreted in the context of how governance controls enter the model through the effective pesticide pressure *ū*_eff_ (*x*_*u*_, *f*). Importantly, because governance enters the system through the aggregated effective pesticide pressure *ū*_eff_ (*x*_*u*_, *f*), the two control pathways are not structurally symmetric. Policy acts as a multiplicative scaling on total pesticide use, whereas adoption redistributes pesticide intensity between management types. As a result, part of the stronger response to policy observed in the (*x*_*u*_, *f*) plane arises from this geometric asymmetry in how controls map to pesticide pressure, rather than from purely ecological differences (Sec. C in Supporting Information).

In the analysis, we use *u*_*H*_ = 1.0 and *u*_*L*_ = 0.5. This choice places the baseline model at the threshold ratio 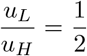 (Sec. C in Supporting Information), where the control aggregation does not impose a uniform 2 built-in marginal advantage of policy over adoption throughout the full (*x*_*u*_, *f*) domain. To define interaction strengths from the binary empirical network, we use degree-scaled couplings

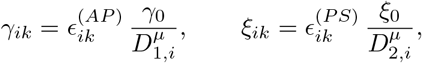

where 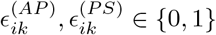 indicate the presence of pollinator-plant and plant-pest links, *D*_1,*i*_ and *D*_2,*i*_ are the corresponding node degrees, and *µ* is a trade-off exponent. This scaling prevents highly connected species from receiving unrealistically large per-link interaction strengths. Parameter values used in the baseline simulations are summarized in Table 2.

**Table 1:** Baseline parameter values used in the tripartite P4 model.

| Symbol | Meaning | Value |
| --- | --- | --- |
| $\alpha_i^{(A)}, \alpha_i^{(P)}, \alpha_i^{(S)}$ | Intrinsic growth rates | 0.1 |
| $\kappa_i^{(A)}, \kappa_i^{(S)}$ | Background mortality rates | 0.0001 |
| $h$ | Handling-time parameter in the saturating responses | 0.7 |
| $m_{A,i}$ | Pesticide sensitivity of pollinator $i$ | 0.5 |
| $m_{S,i}$ | Pesticide sensitivity of pest $i$ | 0.9 |
| $\gamma_0$ | Baseline mutualistic coupling strength | 1.8 |
| $\xi_0$ | Baseline antagonistic coupling strength | 1.0 |
| $u_H, u_L$ | Conventional and IPM pesticide intensities | 1.0, 0.5 |
| $x_u$ | Fraction of farmers adopting IPM | $[0, 1]$ |
| $f$ | Policy control parameter | $[0, 1]$ |
| $\beta_{ii}^{(A)}, \beta_{ii}^{(P)}, \beta_{ii}^{(S)}$ | Intraspecific competition coefficients | 1.0 |
| $\beta_{ij}^{(A)}, \beta_{ij}^{(P)}, \beta_{ij}^{(S)}$ | Interspecific competition coefficients | 0.1 |
| $\mu$ | Degree-strength trade-off exponent | 0.5 |

**Table 2:** Two-hop exposure metrics for pollinators in the empirical tripartite network. For each pollinator *a, k*_*A*_ denotes pollinator degree, *L* is the total two-hop pest burden, 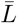 is the degree-normalized burden, *L*_max_ is the maximum pest degree among neighbouring plants, and *L*_2_ is the quadratic burden. The final column records recovery status under the reference management setting, where R and NR denote recovered and non-recovered pollinators, respectively.

| Pollinator | $k_A$ | $L$ | $\bar{L}$ | $L_{\max}$ | $L_2$ | R/NR |
| --- | --- | --- | --- | --- | --- | --- |
| 1 | 1 | 19 | 19.0 | 19 | 361 | NR |
| 2 | 1 | 19 | 19.0 | 19 | 361 | NR |
| 3 | 1 | 0 | 0.0 | 0 | 0 | R |
| 4 | 1 | 1 | 1.0 | 1 | 1 | NR |
| 5 | 3 | 1 | 0.333 | 1 | 1 | R |
| 6 | 2 | 1 | 0.5 | 1 | 1 | NR |
| 7 | 2 | 1 | 0.5 | 1 | 1 | R |
| 8 | 2 | 5 | 2.5 | 5 | 25 | NR |
| 9 | 1 | 5 | 5.0 | 5 | 25 | NR |
| 10 | 1 | 0 | 0.0 | 0 | 0 | NR |
| 11 | 1 | 0 | 0.0 | 0 | 0 | R |
| 12 | 1 | 1 | 1.0 | 1 | 1 | R |
| 13 | 1 | 1 | 1.0 | 1 | 1 | NR |
| 14 | 3 | 24 | 8.0 | 19 | 386 | NR |
| 15 | 1 | 0 | 0.0 | 0 | 0 | NR |
| 16 | 1 | 19 | 19.0 | 19 | 361 | NR |
| 17 | 3 | 2 | 0.667 | 1 | 2 | R |
| 18 | 7 | 3 | 0.429 | 1 | 3 | R |
| 19 | 1 | 1 | 1.0 | 1 | 1 | R |
| 20 | 1 | 5 | 5.0 | 5 | 25 | NR |
| 21 | 2 | 1 | 0.5 | 1 | 1 | R |
| 22 | 1 | 0 | 0.0 | 0 | 0 | NR |
| 23 | 1 | 5 | 5.0 | 5 | 25 | NR |

### 4.4 Synthetic networks with controlled modularity

To isolate the effect of network architecture, we generated 30 synthetic tripartite networks in which only the pollinator-plant layer was rewired. The plant-pest layer, species richness in each guild, and the degree sequence of the pollinator-plant layer were held fixed. Rewiring therefore changes the modular organization of the mutualistic layer without altering species richness, specialization level, or the antagonistic layer. Recovery thresholds and recovered fractions were then recomputed in each synthetic network using the same dynamical protocol as in the empirical network and related to the resulting pollinator-plant modularity (Sec. B.2 in Supporting Information).

### 4.5 Basin stability and initial-condition sensitivity

For the three-dimensional system with state variables (*A, P, S*), the basin stability [48, 49] of a basin of attraction ℬ is defined as

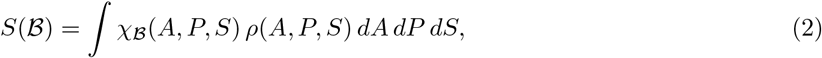

where *ρ*(*A, P, S*) denotes the distribution of initial conditions and *χ*ℬ is the indicator function of the basin of attraction ℬ,

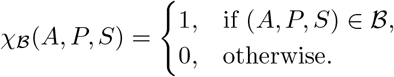

For numerical estimation, we sample *M* initial conditions from a bounded admissible set *I* ⊂ ℝ^3^, integrate the dynamics forward in time, and classify outcomes by their long-time behaviour (Sec. A in Supporting Information). An equilibrium 𝒜 is asymptotically stable if the state variable satisfy

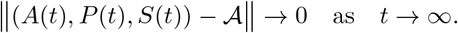

We draw *M* initial conditions independently from *I*, integrate the dynamics forward in time, and record whether trajectories converge to 𝒜. If *N* out of the *M* trials converge to 𝒜, the basin stability of 𝒜 is estimated as

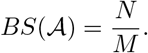

In this study, using this BS approach, two asymptotically stable states of the pollinator with the highest degree node of the (*N*_*A*_ + *N*_*P*_ + *N*_*S*_)-dimensional model (Eq. 1) under the tripartite setup are evaluated, corresponding to extinct (zero) and non-extinct (non-zero) states. For simplicity, the (*N*_*A*_ + *N*_*P*_ + *N*_*S*_)-dimensional state space is reduced to three dimensions by initializing all nodes within each guild with identical abundances, i.e. 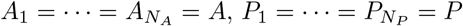, and 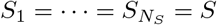. For the calculation of BS, initial conditions are collected uniformly on a 30 ×30 × 30 grid over *A, P, S* ∈ (0, 1], yielding *M* = 27,000 initial conditions. Each sampled point is integrated forward under Eq. 1, and the trajectory is classified according to whether it converges to persistence or extinction of the pollinator. For Fig. 3a, BS of non-extinct state is computed for the highest-degree pollinator (*A*_18_). This approach provides the one-parameter BS curves in Fig. 3b,c.

## Author contributions

A.D. and A.R. conceptualized the research and defined the problem; A.D. and A.R. carried out the implementation. A.R. and A.D. performed data analysis, and A.R, A.D, and U.B. evaluated the results. A.D. and A.R. drafted the initial manuscript. U.B. contributed through revision, editing, feedback, and supervision. A.D., A.R., and U.B. collaboratively finalized the manuscript.

## Acknowledgement

We acknowledge primary funding support from IIT Gandhinagar. The authors acknowledge the ANRF (SERB) Network of networks grant (awarded to U.B) RES/SERB/CE/P0291/2324/0044. U.B. also acknowledges support from the Pandya Shivpuri Chair in Population Dynamics. The authors also extend thanks to H. Poonia, A. S. Kumar, and A. Borah of the Machine Intelligence and Resilience Laboratory at IIT Gandhinagar for their valuable discussions and constructive feedback on this manuscript. A.R. is grateful to S. Ghosh, Postdoctoral Researcher at Lodz University of Technology, for valuable discussions.

## Supporting Information

### A Common numerical conventions

All analyses use the observed tripartite network [1], and the baseline P4 model (Eq. 1 in the main manuscript). The pollinator-plant and plant-pest layers are treated as binary adjacency matrices, while dynamical interaction strengths are constructed from these adjacencies using the same degree-scaled parameterization. Numerical integrations are performed with a fixed-step fourth-order Runge-Kutta algorithm with step size 0.01. All figures are generated from the numerical simulations of the model over 10^5^ iterations.

For the numerical estimation of the basin of attraction associated with the extinction state, each simulated trajectory is classified according to its long-time pollinator abundance. A trajectory is considered to approach the extinction state if

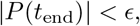

where *t*_end_ is the final integration time. For this study, we fix the value of *ϵ* at 0.01. Trajectories satisfying

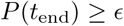

are classified as persisting (non-extinct) states.

### B Network structural diagnostics of recovery heterogeneity and robustness analyses

The same pollinator degree does not imply that the pollinator becomes recovered by varying management parameters because degree counts only the number of plant neighbors and ignores how those neighbors are embedded in the pest layer. Two pollinators may each have degree *k*_*A*_ = *d* (say), yet one may be connected to plants with weak pest exposure while the other is connected to plants that are strongly coupled to multiple pests. As a consequence, they experience different indirect pest burdens despite having identical direct connectivity. Figure 5 depicts that the same pollinator degree does not necessarily imply the same recovery outcome. For instance, pollinator nodes 12 and 9 both have degree *k*_*A*_ = 1, indicating that each is connected to only a single plant. However, their recovery outcomes differ because the neighboring plants host substantially different pest abundances. Pollinator 12 is associated with a plant experiencing relatively low pest pressure, whereas pollinator 9 is connected to a plant that is more strongly linked to pest populations. Consequently, despite having identical direct degrees, the two pollinators are subjected to markedly different indirect pest pressures. As a result, under the influence of the management parameter, node 12 recovers while node 9 does not. A similar contrast appears for pollinator nodes 7 and 8, both of which have degree *k*_*A*_ = 2. Despite having the same number of plant partners, pollinator 7 is embedded in a much less pest-exposed two-hop neighborhood than pollinator 8. The motif snapshots therefore provide a direct structural explanation for why pollinators with the same immediate connectivity can exhibit different recovery behavior under the same management setting. These examples demonstrate that recovery is not determined by the number of plant neighbors alone. Instead, it also depends on how strongly those neighboring plants are themselves embedded in the pest layer. This mechanism will be further clarified in the following subsection using network-based features.

**Figure 5:**
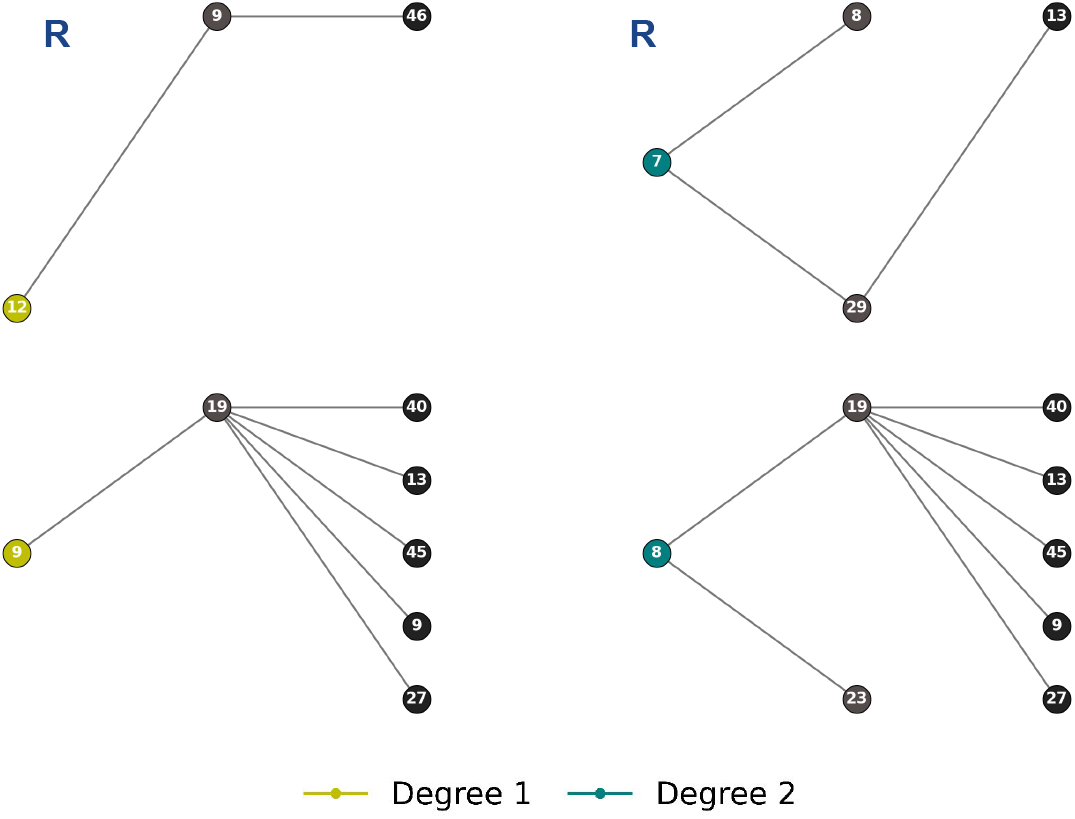
Motif for low-degree pollinators in the empirical tripartite network. Pollinators of degree 1 and 2 are shown together with their neighboring plant and pest nodes. The label R marks recovered pollinators under the reference management setting, whereas unlabeled nodes denote non-recovered pollinators.

#### B.1 Two-hop structural diagnostics of recovery heterogeneity

To understand why pollinators with the same degree can nevertheless display different recovery outcomes under identical stress conditions (as discussed above), we move beyond pollinator connection with plants (pollinator degree) and examine their two-hop structural environment. In the empirical tripartite network, the pollinator degree *k*_*A*_ counts only the number of plant neighbours of a pollinator and therefore provides a strictly local description of topology. By construction, it cannot distinguish whether those plant neighbours are themselves weakly or strongly exposed to the pest layer [2]. Consequently, degree alone is not sufficient to explain why pollinators with the same *k*_*A*_ may bifurcate into different recovery states. To resolve this limitation, we introduce a simple two-hop description that quantifies how strongly each pollinator is indirectly embedded in the pest layer through its neighbouring plants.

Let *N* (*a*) denote the set of plant neighbours of pollinator *a*, and let *k*_*S*_(*p*) denote the pest degree of plant *p*, that is, the number of pest neighbours attached to that plant. We define the following two-hop descriptors.

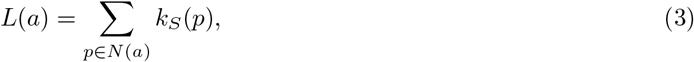

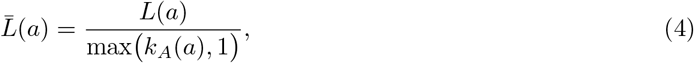

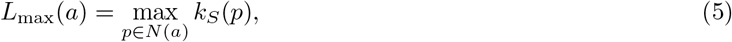

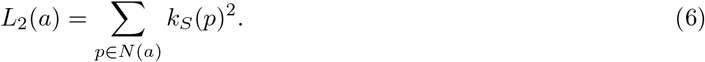

Here, *L*(*a*) measures the total pest burden transmitted through a pollinator’s neighbouring plants, 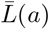 normalizes this burden by pollinator degree, *L*_max_(*a*) captures the worst local exposure among its plant partners, and *L*_2_(*a*) amplifies concentration of burden by weighting highly pest-connected plants more strongly. These quantities depend only on the binary tripartite adjacency structure and are intended to capture a minimal structural mechanism through which equal degree can still yield unequal recovery outcome.

Two pollinators may have the same number of plant links, yet one may be connected to plants that are heavily shared with pests, whereas the other may be connected to plants that are comparatively insulated from pest pressure. Such differences are invisible to *k*_*A*_ alone, but they are directly encoded by the two-hop quantities in Eqs. (3)-(6). For each pollinator node, these metrics were calculated directly from the empirical adjacency structure (Table 2), and each node was labeled as recovered (R) or non-recovered (NR) according to the dynamical outcome of the P4 model under the reference management pamaeters.

Figure 6 presents a descriptive boxplot comparison of these motif-derived quantities between recovered and non-recovered pollinators. The top panel accumulates pollinator nodes by each degree class (*k*_*A*_ = 1, 2, 3, 7) and, within each class, compares the distributions of 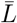, *L*_max_, and *L*_2_ between the two outcome classes. The bottom panel brings all degree classes and repeats the same comparison at the network level. Across all three metrics, non-recovered pollinators tend to occupy more pest-exposed two-hop environments than recovered pollinators. This pattern is exhibited in the comparisons, where the NR class is shifted upward relative to the R class for 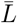, *L*_max_, and *L*_2_. Thus, recovery heterogeneity is not only explained by how many plant partners a pollinator has, but also by how strongly those partners are themselves coupled to the pest layer.

**Figure 6:**
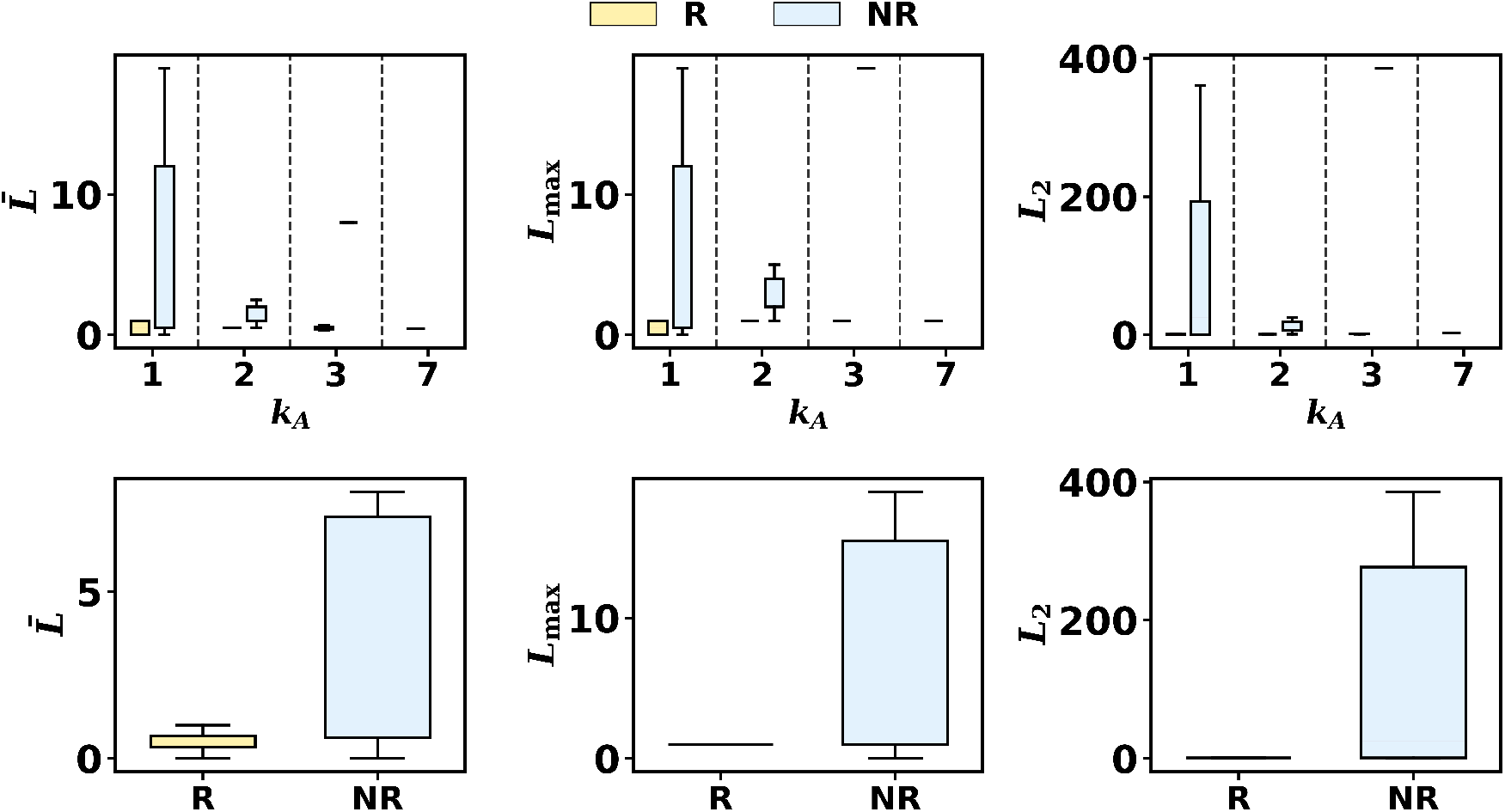
Degree-resolved and pooled comparisons of two-hop motif metrics between recovered and non-recovered pollinators. Top row: boxplots of the degree-normalized burden 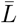, the worst-case burden *L*_max_, and the quadratic burden *L*_2_ across pollinator degree classes. Bottom row: pooled comparisons across all degrees. In each case, non-recovered pollinators tend to exhibit larger two-hop pest-exposure metrics than recovered pollinators, indicating stronger indirect embedding in the pest layer.

Figure. 6 also makes clear that two-hop structure is not a complete determinant of outcome. The recovered and non-recovered classes are not perfectly separated, and there is visible overlap in the distributions (*k*_*A*_ = 1, *k*_*A*_ = 2). This indicates that although higher two-hop pest burden biases a pollinator towards non-recovery, similar local structural environments can still lead to different dynamical outcomes. In other words, the motif variables capture an important structural tendency, but not the full state-dependent nonlinear response of the tripartite ecological system.

To assess whether two-hop structural descriptors provide predictive information beyond direct pollinator degree, we formulate recovery prediction as a binary classification problem. Let *y*_*a*_ ∈ {0, 1} denote the recovery outcome of pollinator *a*, where *y*_*a*_ = 1 indicates recovery (*R*) and *y*_*a*_ = 0 denotes non-recovery (*NR*) under the reference management setting. For each pollinator, we construct feature vectors based on network topology. The baseline representation is given by the scalar feature *x*_*a*_ = *k*_*A*_(*a*), where *k*_*A*_(*a*) denotes the pollinator degree. Motif-aware representations augment this with one two-hop descriptor, yielding feature sets of the form 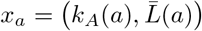, *x*_*a*_ =(*k*_*A*_(*a*), *L*_max_(*a*)), or *x*_*a*_ = (*k*_*A*_(*a*), *L*_2_(*a*)).

For each feature set, we fit a logistic regression model of the form [3]

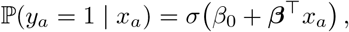

where *σ*(*z*) = (1 + *e*^−*z*^)^−1^ is the logistic link function, and (*β*_0_, ***β***) are estimated via maximum likelihood. Model performance is evaluated using repeated stratified *K*-fold cross-validation, which preserves the class proportions of recovered and non-recovered pollinators within each fold. For each split, the model is trained on the in-fold data and evaluated on the held-out subset.

Predictive performance is quantified using the area under the receiver operating characteristic curve (AUC) [4],

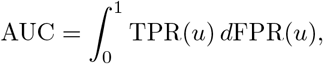

which measures the probability that the classifier assigns a higher score to a randomly chosen recovered pollinator than to a non-recovered one. For each model, the mean AUC across all cross-validation realizations is reported, and the corresponding variability is represented by error bars.

Figure 7 summarizes these results by comparing the baseline (degree-only) model with motif-augmented models. This framework provides a controlled statistical test of whether incorporating two-hop pest-exposure descriptors enhances discriminative power beyond immediate connectivity, thereby quantifying the predictive contribution of higher-order network structure to recovery outcomes.

**Figure 7:**
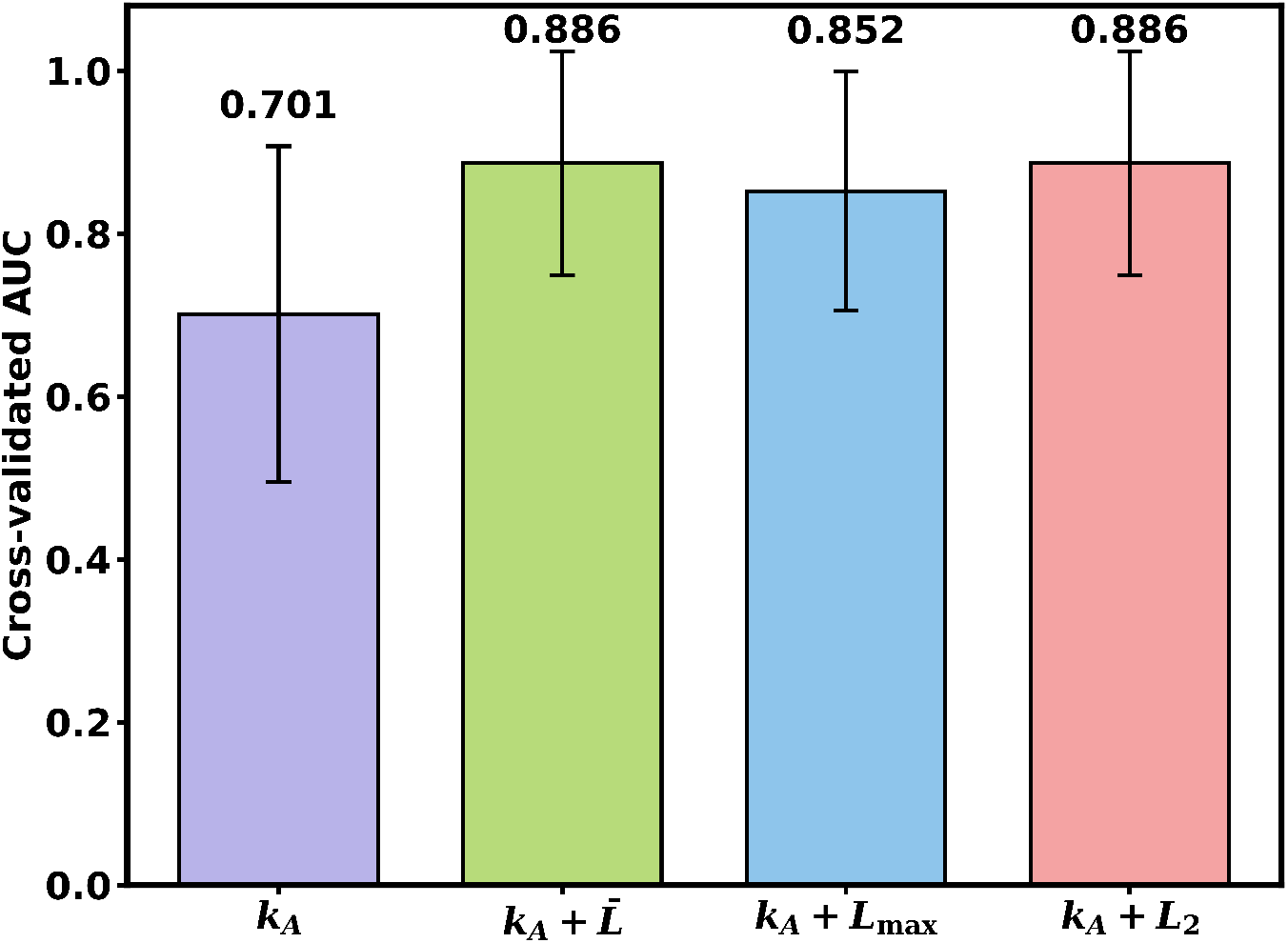
Cross-validated classifier performance for degree-only and motif-aware predictors. Bars show the mean cross-validated AUC for logistic classifiers using *k*_*A*_ alone and *k*_*A*_ augmented with 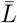, *L*_max_, or *L*_2_. Error bars denote variability across repeated stratified cross-validation folds. The improved performance of motif-aware models shows that two-hop structural descriptors provide predictive information beyond direct pollinator degree alone.

These gains show that two-hop structural information carries predictive content that is absent from direct degree. A pollinator’s recovery therefore depends not only on the number of plant neighbours it possesses, but also on the pest-load architecture of those neighbours. However, the predictive test again supports a qualified rather than absolute interpretation. Although motif-aware models substantially outperform the degree-only baseline, none reaches perfect discrimination, and the permutation-based comparisons with the baseline remain suggestive rather than definitive at conventional thresholds (*p* ≈ 0.099 for 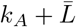, *p* ≈ 0.188 for *k*_*A*_ + *L*_max_, and *p*≈ 0.089 for *k*_*A*_ + *L*_2_). Together with the overlap seen in Fig. 6, this indicates that two-hop exposure is an important predictor of recovery propensity, but not a complete description of recovery outcome.

Figures. 6 and 7 support a common mechanistic picture. The descriptive analysis shows that non-recovered pollinators are, on average, embedded in more pest-loaded two-hop neighbourhoods, while the predictive analysis shows that incorporating such motif-aware information improves discrimination beyond degree alone. Recovery is not governed by topology alone, but by the interplay between topology and nonlinear dynamics, indirect feedbacks among plants, pollinators, and pests, heterogeneity in interaction and the collective network state all influence whether a given pollinator ultimately recovers. In this sense, two-hop pest burden is best interpreted as a statistically informative and mechanistically meaningful contributor to recovery heterogeneity, rather than as a single sufficient cause.

#### B.2 Structural effects of modularity on pollinator recovery

To examine whether recovery heterogeneity is shaped not only by local two-hop exposure but also by higher-order network architecture, we next analyzed a controlled ensemble of synthetic tripartite networks with varying pollinator-plant modularity. The goal here is to isolate the contribution of large-scale structural organization to recovery thresholds and recovery extent. Whereas the previous analysis showed that pollinators with the same degree can experience different outcomes because of differences in their immediate two-hop pest environment, the present analysis asks whether the mesoscopic arrangement of the mutualistic layer itself systematically biases the recovery landscape.

We have generated 30 synthetic pollinator-plant-pest networks in which only the pollinator-plant layer was rewired to achieve different levels of modularity, while the plant-pest layer, species richness of each guild, and the degree sequence of the pollinator-plant layer are kept fixed. This design ensures that any observed differences in recovery arise primarily from changes in the compartmental organization of the mutualistic layer rather than from changes in richness, connectance, or antagonistic structure. For each synthetic network, we then calculate the pollinator-plant modularity and evaluated the recovery dynamics using the same dynamical protocol as in the empirical network for two different management scenario.

Figure 8 shows that recovery outcomes depend strongly on this architectural variation. As modularity increases, the critical value of IPM adoption 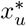 at fixed policy intensity *f* = 0.1 increases markedly (Fig. 8a), indicating that more modular mutualistic organization requires a larger level of external intervention before recovery can occur. A similarly increasing trend is observed for the critical value of policy *f*^*^ at fixed *x*_*u*_ = 0.1 (Fig. 8c), showing that stronger external intervention is needed as the network becomes more compartmentalized. In parallel, the fraction of pollinator species that successfully transition to recovery declines with modularity both at 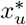 and at *f*^*^ (Fig. 8b, d).

**Figure 8:**
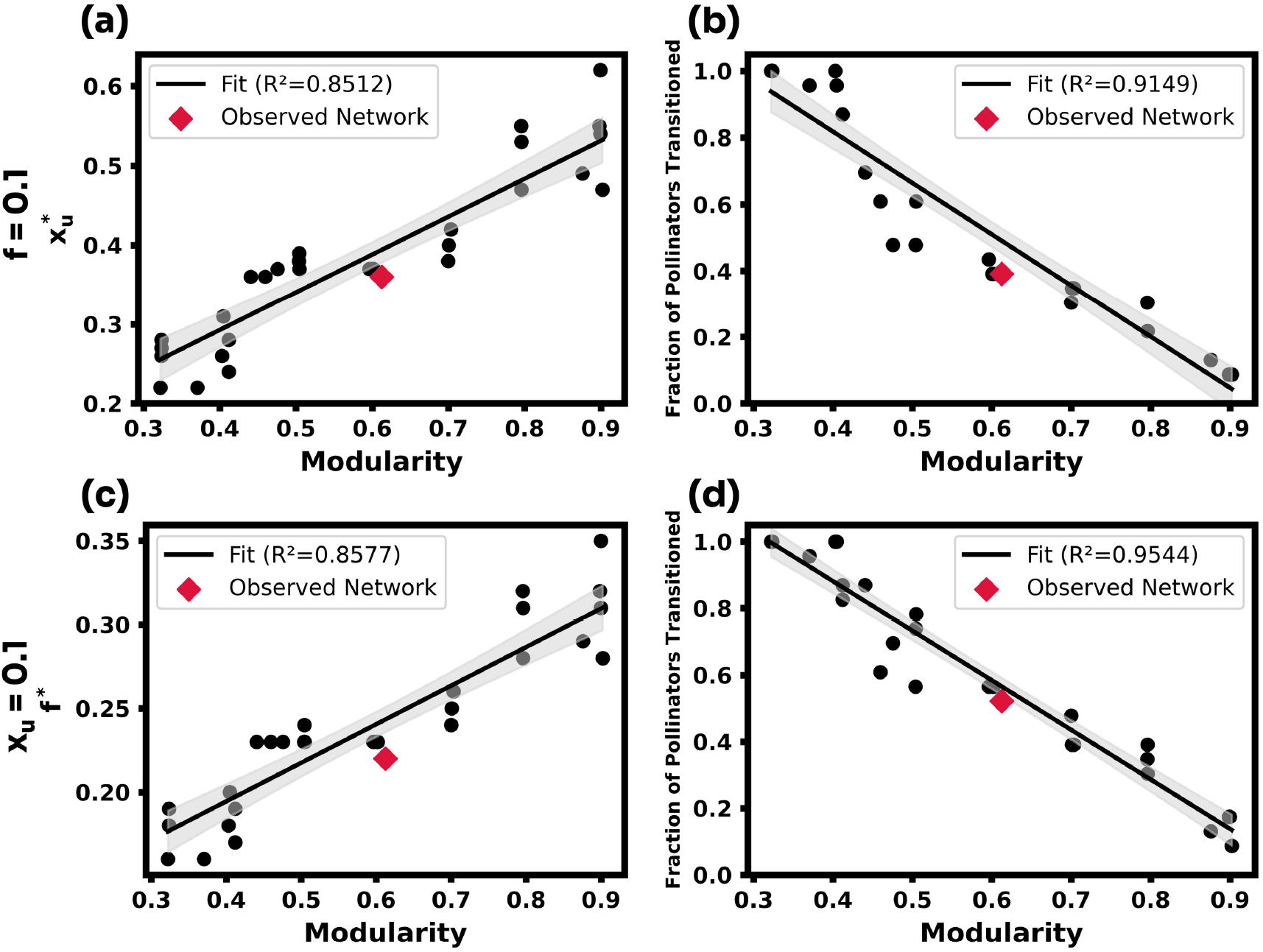
Effect of pollinator-plant modularity on recovery thresholds and recovery extent across 30 synthetic tripartite networks. (a) Critical adoption threshold 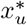 at fixed *f* = 0.1. (b) Fraction of pollinator species that recover at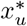. (c) Critical policy threshold *f*^*^ at fixed *x*_*u*_ = 0.1. (d) Fraction of pollinator species that recover at *f*^*^. Points denote individual synthetic networks, solid lines show ordinary least-squares fits with 95% confidence bands, and red diamonds mark the empirical network. Increasing modularity raises recovery thresholds and reduces the fraction of recovered pollinators.

These fitted trends explain a substantial fraction of the variation across synthetic networks, with *R*^2^ = 0.8512 for 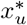 versus modularity, *R*^2^ = 0.9149 for the recovered fraction at 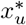, *R*^2^ = 0.8577 for *f*^*^ versus modularity, and *R*^2^ = 0.9544 for the recovered fraction at *f*^*^. The empirical network, marked separately in Fig. 8, lies close to these fitted relationships, indicating that it follows the same structural tendency observed across the synthetic ensemble. This agreement supports the interpretation that the modular organization of the pollinator-plant layer is itself a meaningful determinant of recovery behavior.

The result suggests that stronger modularity restricts the propagation of beneficial recovery effects across nodes. In a weakly modular network, recovery initiated in one part of the pollinator-plant layer can more readily spread through cross-module connections, allowing persistence gains to percolate across the system. By contrast, in a highly modular network, interactions are more compartmentalized, so recovery remains more localized and requires stronger management forcing to cross structural boundaries.

### C Disentangling control leverage from ecological dynamics through matched-stress comparison

In the main framework of the proposed dynamical system (Sec. 4.2 and Sec. 4.3 in the main manuscript), pesticide governance enters the tripartite ecological system through the effective pesticide pressure

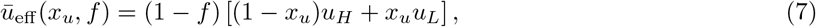

where *f* ∈ [0, 1] denotes the strength of centralized policy intervention, *x*_*u*_ ∈ [0, 1] denotes the fraction of decentralized IPM adoption, *u*_*H*_ and *u*_*L*_ are the pesticide intensities associated with conventional and IPM management. This representation combines both governance pathways into a single scalar pressure term that acts on the ecological dynamics. However, it also implies that the two control parameters do not enter symmetrically. Consequently, a direct comparison between policy and adoption in the (*x*_*u*_, *f*) parameter plane may conflate two distinct effects, namely the intrinsic nonlinear ecological response of the system and the geometric asymmetry of the control map itself.

To expose this asymmetry [5], we consider that the local (marginal) leverage on pesticide pressure is governed by the partial derivatives of Eq. (7). Differentiating partially yields

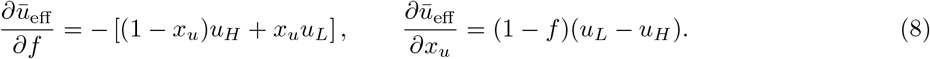

Taking magnitudes gives

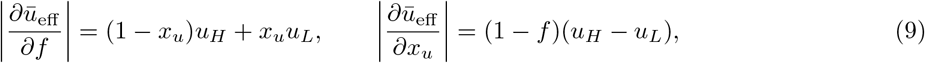

so that the relative marginal influence of *f* and *x*_*u*_ varies over (*x*_*u*_, *f*). In particular, the two parameters are not locally symmetric under the multiplicative coupling in Eq. (7) because 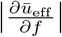 depends on *x*_*u*_ but not on *f*, whereas 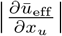 depends on *f* but not on *x*_*u*_, i.e. the marginal leverage of policy depends on the current adoption state *x*_*u*_, whereas the marginal leverage of adoption depends on the current policy state *f*. Therefore, a larger apparent gain from policy in direct parameter-space comparisons need not arise solely from ecological superiority, part of it may be induced by the algebraic structure of Eq. (7). So, a structural marginal policy advantage is obtained when

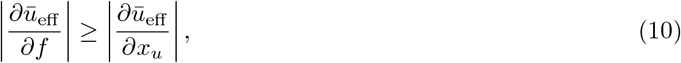

which implies

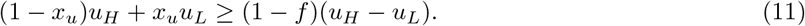

Since *ū* (*x*_*u*_) is linear equation in *x*_*u*_ and decreases from *u*_*H*_ at *x*_*u*_ = 0 to *u*_*L*_ at *x*_*u*_ = 1, we have 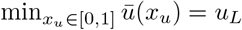. Also, (1 − *f*)(*u*_*H*_ − *u*_*L*_) is maximized at *f* = 0, giving 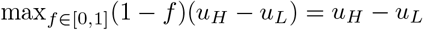. Consequently, Eq. (10) holds for all (*x*_*u*_, *f*) ∈ [0, 1] *×* [0, 1] if and only if

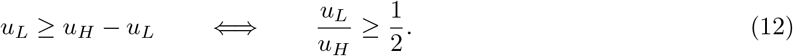

When

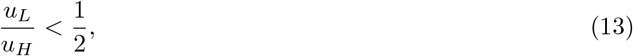

there exist regions in control space where adoption can have greater local leverage than policy. Rearranging Eq. (11), the regime in which adoption dominates locally is

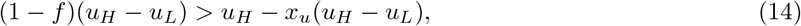

or equivalently,

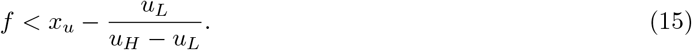

These relations clarify that the control parameters are not directly comparable in a consistent manner unless one accounts for the achieved reduction in effective pesticide burden.

To isolate the ecological response from this structural asymmetry, we therefore construct a comparison in which policy and adoption are constrained to achieve the same effective pesticide pressure. Let (*x*_0_, *f*_0_) be a fixed baseline point and define the baseline effective pesticide level

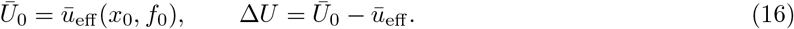

Here, Δ*U* measures the reduction achieved in effective pesticide pressure relative to the baseline state. For policy-only variation, the adoption is kept fixed at *x*_*u*_ = *x*_0_, and the target state (*x*_0_, *f*_new_) is defined implicitly through

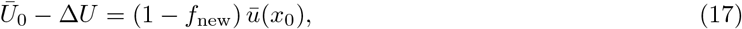

Where

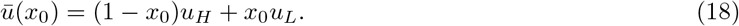

Solving for *f*_new_ gives

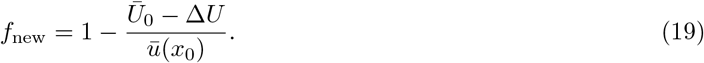

Similarly, for adoption-only variation, policy is held fixed at *f* = *f*_0_, and the matched state (*x*_*u*,new_, *f*_0_) satisfies

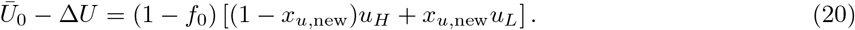

This yields

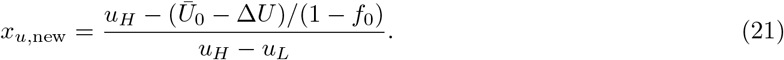

Equations (19) and (21) guarantee that policy and adoption are compared only after enforcing the same achieved decrease in effective pesticide pressure. In this way, the comparison is performed at equal scalar stress reduction rather than at the same parameter displacement. For the numerical experiments, we fix the baseline parameters at (*x*_0_, *f*_0_) = (0.1, 0.1) and set *u*_*H*_ = 1.

The results shown in Fig. 9 are obtained by systematically constructing matched control trajectories using Eqs. (19) and (21), and evaluating the resulting ecological dynamics through numerical integration of the full tripartite dynamical system. For each value of the normalized stress reduction 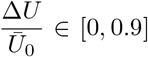 the corresponding policy-controlled and adoption-controlled states are calculated such that both produce identical effective pesticide pressures. The Fig. 9 depicts the change in basin stability (BS) (Sec. 4.5 in the main manuscript), each in three reduced-pesticide values *u*_*L*_ = 0.2, 0.5, and 0.8. The BS for the high degree pollinator (here *A*_18_) is estimated by sampling an 20 *×* 20 *×* 20 grid of initial conditions over (*A, P, S*) ∈ [0.001, 1] *×* [0.001, 1] *×* [0.001, 1].

**Figure 9:**
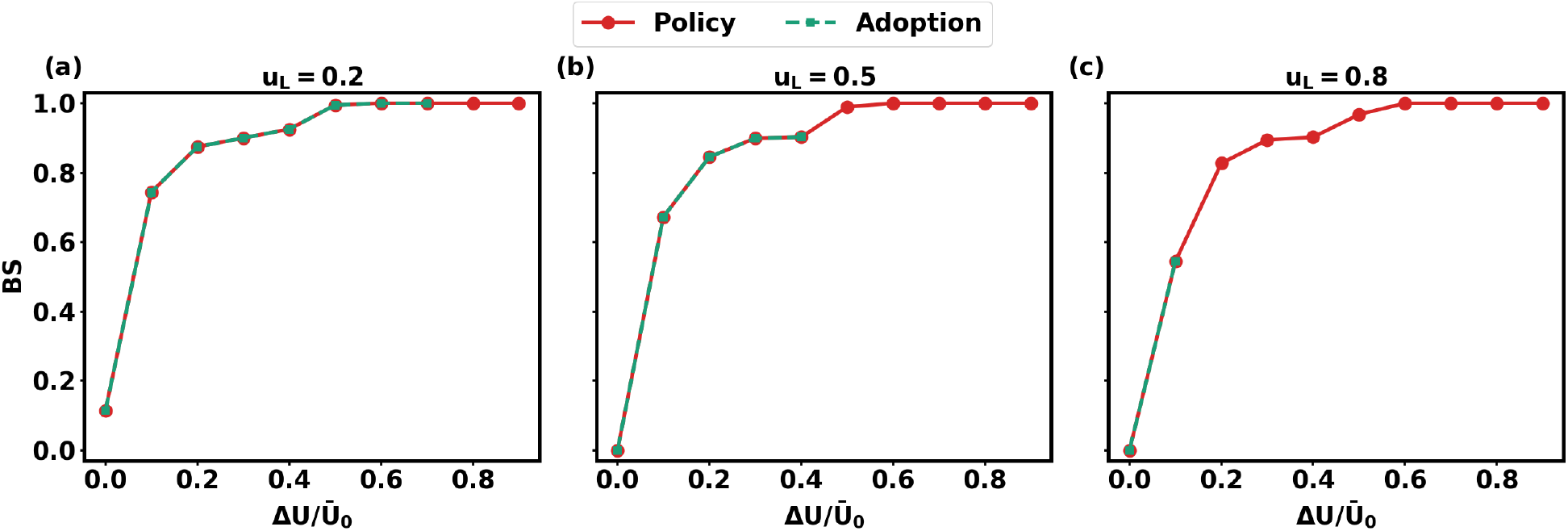
Identical-stress comparison between policy and adoption control pathways across three residual pesticide levels, *u*_*L*_ = 0.2, 0.5, and 0.8. This figure shows basin stability (BS) as a function of the normalized reduction in effective pesticide pressure, Δ*U/Ū*_0_, for policy-controlled (red) and adoption-controlled (green) trajectories, both constrained to achieve the same target value of *ū*_eff_. In each column, policy (*f*_*new*_) varies at fixed *x*_*u*_ = *x*_0_ = 0.1, whereas IPM adoption (*x*_*u,new*_) varies at fixed *f* = *f*_0_ = 0.1. The consistency of this overlap across different *u*_*L*_ values suggests that policy and adoption act as dynamically equivalent mechanisms when constrained to achieve identical effective pesticide pressure.

Under the construction of matched-stress, both the policy and the IPM adoption trajectories satisfy

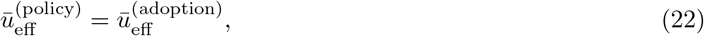

where 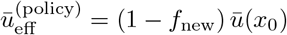,and 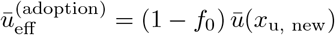,ensuring that the comparison isolates the ecological response under identical environmental forcing. However, the policy and adoption pathways are subject to the feasibility constraint

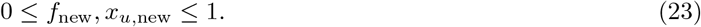

Therefore, for values of Δ*U/Ū*_0_ that yield *f, x*_*u*,new_ *>* 1, the matched policy and adoption state becomes infeasible, and the corresponding points are excluded from the comparison.

Within the feasible domain, Fig. 9 shows that the BS curves for policy (red) and adoption (green) collapse onto a single trajectory, i.e.,

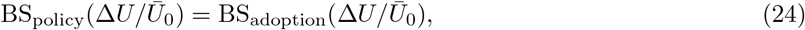

indicating that, within the homogeneous mean-field formulation, the ecological response is determined by the scalar effective pesticide pressure *ū*_eff_. indicating that, within the homogeneous mean-field formulation, the ecological response is determined by the scalar effective pesticide pressure *ū*_eff_. In this regime, the basin stability can be expressed as

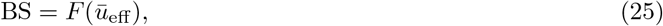

independent of the specific control pathway under the homogeneous mean-field assumption, where *F* be teh arbitrary functional form. This follows because the dynamical system depends on pesticide exposure only through the scalar term *ū*_eff_, so that trajectories corresponding to equal *ū*_eff_ evolve identically.

This formulation implicitly assumes homogeneous exposure, i.e., all species experience the same effective pesticide pressure *ū*_eff_. The monotonic increase of BS with Δ*U/Ū*_0_ reflects a continuous recovery transition, where larger reductions in effective pesticide pressure progressively enlarge the basin of attraction of the recovered state. However, the feasible range of Δ*U/Ū*_0_ depends on the reduced pesticide level *u*_*L*_. As *u*_*L*_ increases, the maximum achievable reduction through adoption becomes limited as *x*_*u,new*_ *>* 1, causing the admissible domain of Δ*U/Ū*_0_ to shrink. Beyond this limit, the adoption pathway becomes infeasible, so further reductions in *ū*_eff_ can only be achieved through policy intervention. Therefore, the matched-stress analysis demonstrates that policy and adoption are dynamically equivalent within the feasible region of the control space. Any apparent advantage of policy arises not from intrinsic ecological differences, but from the broader feasible range of achievable effective pesticide reductions under the policy pathway.

## Notes

https://figshare.com/articles/dataset/Contrasting_effects_of_land-use_changes_on_herbivory_and_pollination_networks/10000352/1

### Competing Interest Statement

The authors have declared no competing interest.

